# deluxpore: a Nextflow pipeline for demultiplexing Illumina dual-indexed Nanopore libraries

**DOI:** 10.64898/2026.03.27.714410

**Authors:** Catalina Arnaiz del Pozo, Claudia Sanchis-López, Jaime Huerta-Cepas

## Abstract

**Summary:** The combination of target capture metagenomics and long-read sequencing represents a powerful approach for the characterisation of rare microbial taxa and their functional genes. However, standard Nanopore library preparations are incompatible with established capture protocols. A possible workaround is the preparation of Illumina libraries prior to ONT sequencing. Currently, this hybrid approach is hindered by a lack of specialised demultiplexing software capable of handling residual adapter fragments; Nanopore’s higher error rates and positional variability. Here, we present deluxpore: a Nextflow pipeline that demultiplexes Nanopore reads from Illumina dual-indexed libraries (NEBNext and Nextera) using BLAST alignment and Levenshtein distance matching. Extensive benchmarking across 18 replicates validates the viability and precision of this hybrid indexing approach. Benchmarking demonstrates that accurate demultiplexing requires minimum Q20 data quality and strategic index selection. Unique index-to-sample designs achieved 91.7% sample recovery at Q20 versus 46.1% for combinatorial approaches. We also identified high-crosstalk index pairs within NEBNext Primer Set A and provide an optimized 8-sample configuration achieving ~95% accuracy at Q20. deluxpore enables reliable, automated demultiplexing for hybrid capture–long-read sequencing workflows.

**Availability and implementation:** ***deluxpore*** is implemented in Nextflow, Python, and Bash under the GNU GPL v3.0. Source code, documentation, and benchmarking workflows are available at https://github.com/compgenomicslab/deluxpore and https://github.com/compgenomicslab/deluxpore-benchmarking.

## 1 Introduction

The rapid advancement of high-throughput sequencing (HTS) has transformed the ability to explore microbial communities, thereby driving demand for scalable and cost-effective solutions (Child et al. 2025). Long-read sequencing, as offered by Oxford Nanopore Technologies (ONT), facilitates the recovery of full-length gene sequences, enabling the functional characterisation of microbial communities to a greater extent than that achievable with short-read technologies (Ciuffreda, Rodríguez-Pérez, and Flores 2021). Nevertheless, both short- and long-read shotgun metagenomics frequently fail to detect rare taxa and low-abundance functional variants, even with extensive sequencing efforts, due to the prevalence of highly abundant organisms within complex communities (Siljanen et al. 2025). This limitation can be addressed by target capture metagenomics, which involves the selective enrichment of preselected genomic regions prior to sequencing (Valmas et al. 2025). This newly prominent approach has been shown to significantly increase the sensitivity of detecting rare microbiota and their associated functional genes (Denonfoux et al. 2013; Manoharan et al. 2015; Jones and Good 2016; Sanchis-López et al. 2025; Kapoor et al. 2026). Therefore, the integration of target capture enrichment with long-read sequencing emerges as a powerful approach for unveiling concealed functional diversity. Probes targeting conserved flanking regions can capture highly divergent gene variants, while long-read sequencing preserves their full-length context for functional characterisation.

However, to fully leverage the power of long-read target capture sequencing across large sample sets, efficient multiplexing strategies are essential. Although ONT barcoding kits facilitate sample multiplexing, they are directly incompatible with established target capture workflows, which are optimized for short-read platforms, making MinION libraries unsuitable for off-the-shelf target enrichment kits (Karamitros and Magiorkinis 2015; Montaguth et al. 2025). As a solution, target enrichment can be performed on Illumina-indexed libraries, whereby dual indexes incorporated during library preparation allow samples to be pooled *prior* to capture hybridisation, after which the enriched pool is converted into a Nanopore-compatible library for MinION sequencing; a strategy recently applied to microbial gene capture using NEBNext Multiplex Oligos (Sanchis-López et al. 2025). Importantly, this hybrid approach lacks practical implementation: standard Illumina demultiplexers cannot process Nanopore-derived data. This is primarily due to the significantly higher raw-read error rates of Nanopore (5–15% vs. 0.1–1% for Illumina), which obscure short index sequences (Murigneux et al., 2021; Bejaoui et al., 2025). Furthermore, traditional algorithms are not designed to recognise indexes embedded within residual adapter fragments, nor are they capable of handling the positional variability typical of long-read sequencing (Renaud et al. 2015; Yi et al. 2015). Consequently, there is a critical need for specialised software capable of robust index identification within high-error, long-read datasets.

The present study proposes a new methodology and software implementation for the demultiplexing of Illumina dual-indexed Nanopore libraries: *deluxpore*, which addresses the compatibility gap between established capture protocols and long-read sequencing. The result is a system that facilitates scalable, high-throughput targeted sequencing on the ONT platform.

## 2 Implementation

### Installation and dependencies

The pipeline requires Conda and Nextflow and supports demultiplexing with NEBNext® Ultra™ II DNA Library Prep Kit (Dual Index Primers Set 1) and Illumina Nextera™ UD Index adapters Set A index libraries. Full installation instructions and parameter options are available in the GitHub repository.

### Method description

Deluxpore is a pipeline that processes Nanopore sequencing reads to demultiplex samples based on Illumina dual-index sequences. Illumina’s dual-index library preparation method employs oligos that feature a conserved adapter sequence, along with a unique index barcode for sample identification. Each oligo contains either an i5 or i7 unique index (8 nt for NEBNext, 10 nt for Nextera), and samples are distinguished by their unique i5/i7 combination. This configuration allows the pipeline to first identify oligo regions via sequence alignment, then extract the adjacent unique index sequences for sample assignment.

The demultiplexing workflow is divided into 4 main stages: (1) **Adapter trimming and quality filtering** to remove Nanopore adapters and low-quality reads; (2) **Adapter sequence identification** through BLAST alignment against a custom database containing only the complete oligo sequences present in the experimental design, reducing computational overhead and spurious mappings; (3) **Unique index extraction and matching**, where unique index sequences are extracted from fixed positions flanking the mapped regions based on unique index length (8 and 10 nt for NEBNext and Nextera, respectively), and compared against the reference unique index library using Levenshtein distance calculations to identify the best-matching i5 and i7 index pair for each read; (4) **Sample assignment** following a hierarchical decision logic.

For each read, the pipeline identifies the best-matching i5 and i7 indexes based on three priority levels: (1) lowest Levenshtein distance to a reference index, (2) alignment position (with i5 indexes prioritised when closer to the read start and i7 indexes when closer to the read end), and (3) validation against the experimental design to ensure the index combination corresponds to a defined sample. In cases where two indexes yield identical distance and position values, but map to different valid sample combinations, the read is flagged as ambiguous and excluded from downstream assignment to prevent misclassification. Therefore, sample assignment is contingent upon the uniqueness of the index within the experimental design. When individual indexes are shared across multiple samples, both i5 and i7 indexes must be successfully identified and matched as a pair for sample assignment (Dual-index assignment). Conversely, in experimental designs where each index uniquely identifies a sample, reads can be assigned based on a single index match (Single-index assignment). This enables recovery of reads where one index failed quality filtering or was not detected. When enabled, the pipeline additionally locates and removes complete oligo sequences from the reads, based on the initial mappings.

### Software implementation

Deluxpore is implemented as a Nextflow pipeline and requires three inputs: (i) Nanopore sequencing reads, either as a single file or divided into chunks; (ii) a parameter file controlling pipeline execution; and (iii) an experimental design file containing tab-delimited sample-to-dual-index associations. Index names must correspond to NEBNext or Nextera unique index sequence identifiers; supported sequences and example input files are documented in the repository.

Reads must be trimmed to remove Nanopore adapters before demultiplexing, either using the pipeline’s built-in Porechop (v0.2.4; https://github.com/rrwick/Porechop) and Chopper (De Coster and Rademakers 2023) modules or pre-processed externally by the user. Trimmed and filtered reads are initially converted to FASTA format using the *seqtk* processing toolkit (https://github.com/lh3/seqtk). Custom Python/Bash scripts handle index extraction, Levenshtein distance calculations, and sample assignment logic.

Computational efficiency is achieved through Nextflow’s parallel execution capabilities to process read chunks concurrently. Details of the assignment and distance metrics are output as JSON files for each chunk, enabling quality assessment, while the demultiplexed reads are concatenated into final per-sample FASTA files. This architecture scales efficiently across both local workstations and HPC environments, with resource allocation configurable through Nextflow profiles.

## 3 Benchmarking

### Dataset

To evaluate the performance of our demultiplexing approach under different error rates and multiplexing scenarios, and establish quality thresholds for high-precision demultiplexing, we created two experimental benchmark datasets simulating realistic sequencing conditions with varying quality levels:

#### 96-sample dataset

The first dataset contained a total of 96 samples, which were multiplexed using a dual-index combinatorial strategy. Theoretical sequences 1,500 nucleotides long were generated using [a custom script](https://github.com/ktlina/deluxpore-benchmarking/blob/main/00-generate_reads.py), where unique combinations of NEBNext dual index Set A i5 and i7 indexes were assigned to each sample. This configuration is representative of a typical high-throughput experimental design, with each sample equimolarly present within the pooled library at 1 million reads per sample. To evaluate the impact of quality scores arising from Nanopore sequencing on demultiplexing performance, the experimental dataset was used as input for 18 independent rounds of BadRead (Wick 2019) Nanopore error simulations at various quality score levels (Q10, Q20, Q25, and Q30). BadRead simulations employed the nanopore2023 error model, with identity values set to 10 ± 3%, 20 ± 3%, 25 ± 3%, and 30 ± 3% for the Q10–Q30 datasets, respectively. The read length was set to 1634 ± 15 bp and artefact generation (adapters, chimeras, junk reads and glitches) was disabled to isolate the effect of sequencing error on demultiplexing performance.

#### 8-sample dataset

The second dataset consisted of 8 samples selected from the 96-sample experimental design. The index combinations were chosen based on the 96-sample design i7 confusion matrix results. One index from each confused pair (i704-i706, i7010-i702 and i7011-i7012) was removed to eliminate cross-assignment errors whilst maximising the number of indexes retained. To enable direct comparison using identical reads, the eight-sample dataset was subsetted from the original 96-sample BadRead simulations rather than being generated independently.

### Demultiplexing Performance

#### Sample assignment rates

Sample assignment rates varied considerably depending on index uniqueness within the experimental design. For the 96-sample dataset, where each index is shared across multiple samples being both indexes required for unambiguous assignment, only 11.8% of reads were successfully assigned at Q10, increasing substantially with higher quality thresholds (**Figure 1**A). In contrast, the 8-sample dataset, where each index uniquely identifies a sample, achieved higher assignment rates: 43.7% at Q10, 91.7% at Q20, 96.1% at Q25, and 97.5% at Q30 (**Figure 1**B). This difference reflects the ability to assign reads based on a single successfully identified index when index-to-sample relationships are unique.

**Figure 1.**
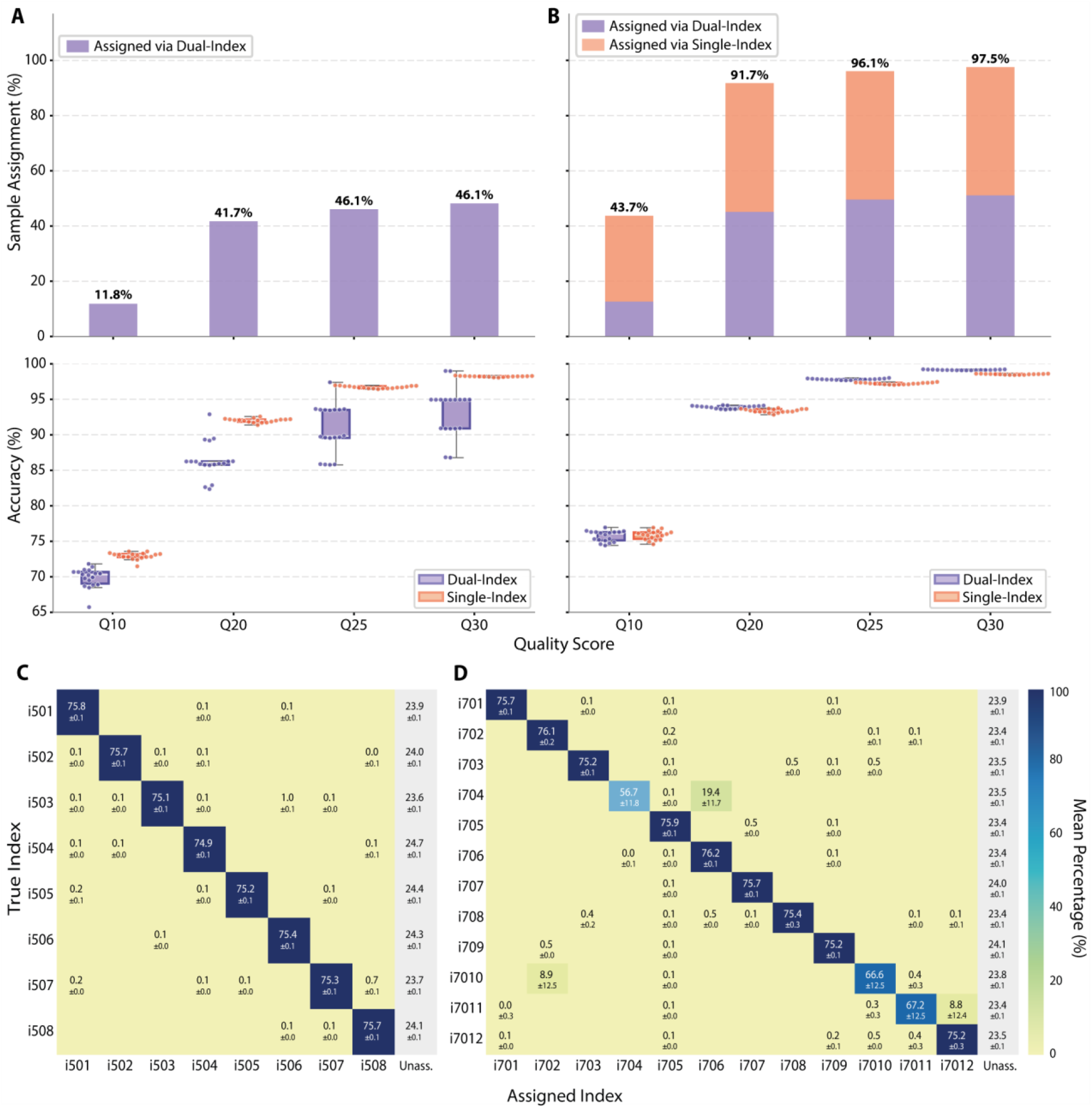
Benchmarking demultiplexing performance across quality score thresholds. **(A, B)** Performance metrics for 96-sample and 8-sample experimental designs, respectively. **Upper panels**: Sample assignment rates showing the proportion of reads successfully assigned to samples. **Lower panels**: Assignment accuracy (%) for dual-index (purple) and single-index (orange) approaches across quality thresholds; dots represent individual replicates (n=18), and lines connect mean values. **(C)** Confusion matrix for the 96-sample dataset showing mean assignment percentages (± std) for i5 indexes (i501-i508) at Q30 quality level. Diagonal values represent the correct assignments; off-diagonal values indicate cross-assignment between indexes. **(D)** Confusion matrix for i7 indexes (i701-i712) from the 96-sample dataset at Q30. Notable confusion pairs (e.g., i704-i706, i7010-i702, and i7011-i7012) exhibit reduced diagonal values and elevated off-diagonal assignments. The “Unassigned” column indicates reads that failed to meet assignment thresholds for confident index calling. The colour scale represents the mean assignment percentage (0–100%).

Notably, at Q25 and above, the 8-sample design achieved near-complete sample assignment (>96%). This design represents a more realistic experimental scenario that aligns with manufacturer recommendations for hybridization capture workflows, which typically suggest pooling eight libraries per capture reaction (https://arborbiosci.com/wp-content/uploads/2022/03/myBaits_v5.02_Manual.pdf). These results demonstrate that while high-throughput dual-index combinatorial designs are feasible, experimental designs employing unique dual-index-to-sample assignments substantially improve read recovery, particularly at lower quality thresholds.

#### Index assignment accuracy

To validate the reliability of index predictions, we assessed assignment accuracy across four quality score thresholds (**Figure 1**A, B, lower panels). In the 96-sample experimental design, dual-index accuracy (purple) remained high but exhibited notable variability at the Q20 and Q25 thresholds, where several replicates showed lower-than-average performance despite the overall upward trend. Conversely, single-index accuracy (orange) demonstrated a more concentrated distribution and a substantial enhancement as quality requirements escalated, rising from around 73% at Q10 to over 98% at Q30.

In contrast, the 8-sample design exhibited a notable improvement in dual-index accuracy and a reduction in replicate variance compared to the 96-sample set. This divergence is attributed to the strategic selection of indexes for the 8-sample subset. By filtering out high-crosstalk indexes, the 8-sample dataset achieved near-perfect accuracy (>98%), even at moderate quality thresholds (Q20). These results confirm that while higher quality thresholds increase confidence in index predictions, the underlying experimental design is a primary driver of demultiplexing reliability.

#### Confusion matrix assignment

To characterize the specific patterns of misassignment, we generated confusion matrices for the 96-sample dataset (**Figure 1**C, D). The i5 indexes (i501–i508) demonstrated almost ideal performance, with mean correct assignment rates consistently exceeding 75% across the diagonal and negligible off-diagonal crosstalk (**Figure 1**C). In contrast, the i7 indexes (i701–i712) revealed specific “hot spots” of significant index confusion (**Figure 1**D). While most performed well, three distinct pairs showed elevated cross-assignment: i7010-i702, i704-i706, and i7011-i7012. This level of confusion was reverted in the 8-sample experimental design, where cross-assignments were reduced to negligible levels (≤0.7%), while correct diagonal assignments remained stable between 75.1% and 75.9% (Supplementary Figure S1).

The “Unassigned” column for both index types accounted for approximately 24% of reads, where the vast majority of reads across all higher quality scores were identified as “Partial” sequences (i5 length: <1565 bp; i7 length: <1568 bp). “Full-length” sequences (≥1634 bp) that failed assignment, represent a minimal fraction of the data, decreasing as quality thresholds increased (Supplementary Figure S2). Therefore, demultiplexing failures in this study are primarily driven by sequence truncation rather than poor base-calling quality of the index regions themselves.

## 4 Conclusions

Our benchmarking results demonstrate that precise demultiplexing for hybridisation capture workflows, coupled with long-read Nanopore sequencing, is achievable by prioritizing sequence-distinct experimental designs alongside sequencing quality. We recommend a minimum data quality of Q20 and the use of unique dual-index-to-sample relationships, as our optimized 8-sample configuration, which excludes high-crosstalk pairs such as i704–i706 and i7011–i7012, achieved >98 % accuracy at Q20. While high-throughput combinatorial designs are technically feasible, they require exceptional quality thresholds (Q30) to match the performance of unique designs at moderate quality. We provide the optimised index combinations and ***deluxpore*** software to enable automated, high-accuracy demultiplexing for hybrid capture–long read workflows.

## Data Availability

The data underlying this article are available in Zenodo, at https://doi.org/10.5281/zenodo.18697344. These datasets include simulated reads and demultiplexing results used for benchmarking.

## SUPPLEMENTARY FIGURES

**Supplementary Figure S1.**
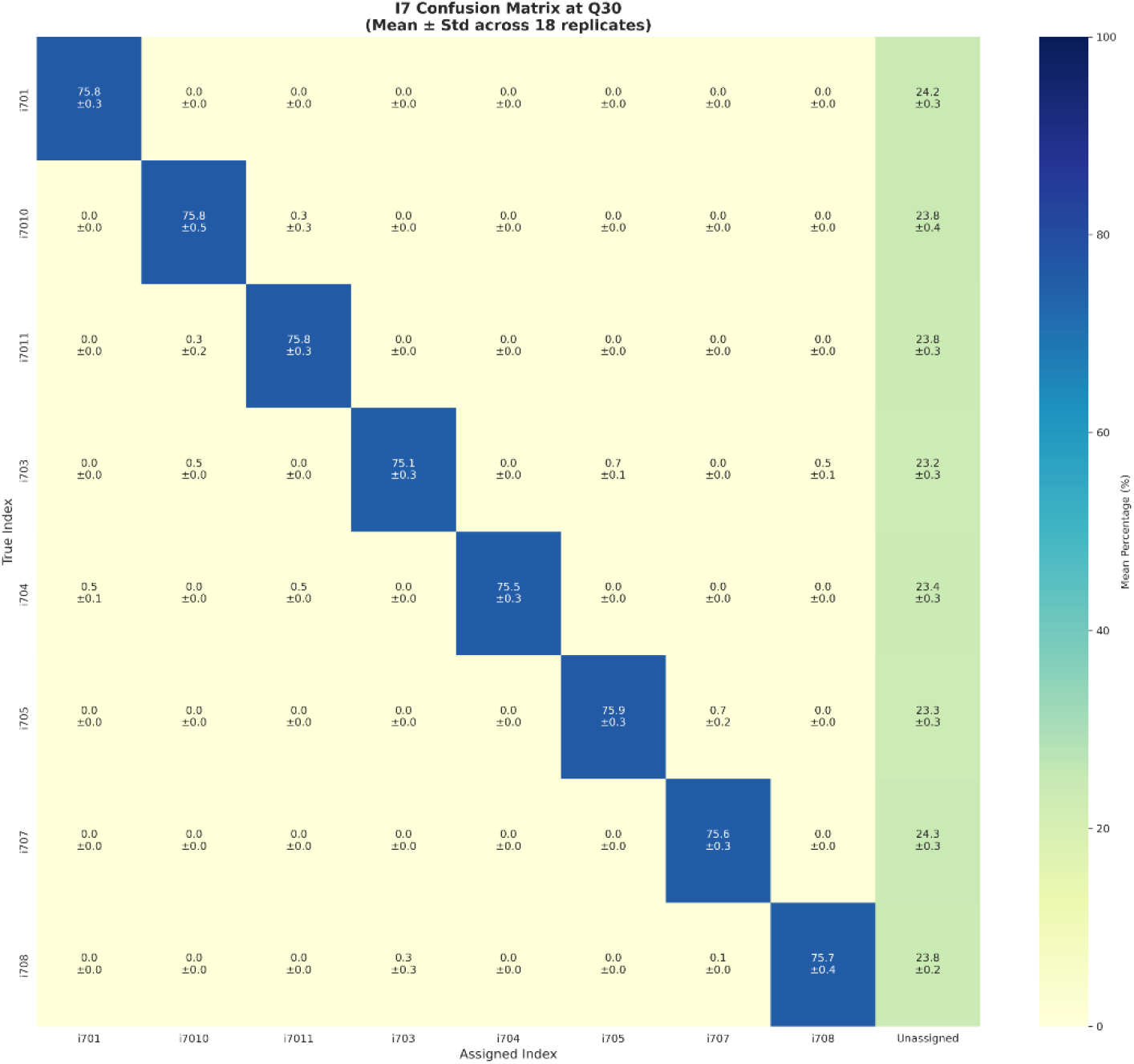
i7 Index Confusion Matrix for the 8-sample dataset. Mean assignment percentages (± standard deviation) are shown across 18 replicates at a Q30 quality threshold. Following the strategic removal of confused index pairs, confusion between indexes is effectively eliminated, with the “Unassigned” column representing the primary reason for non-assignment.

**Supplementary Figure S2.**
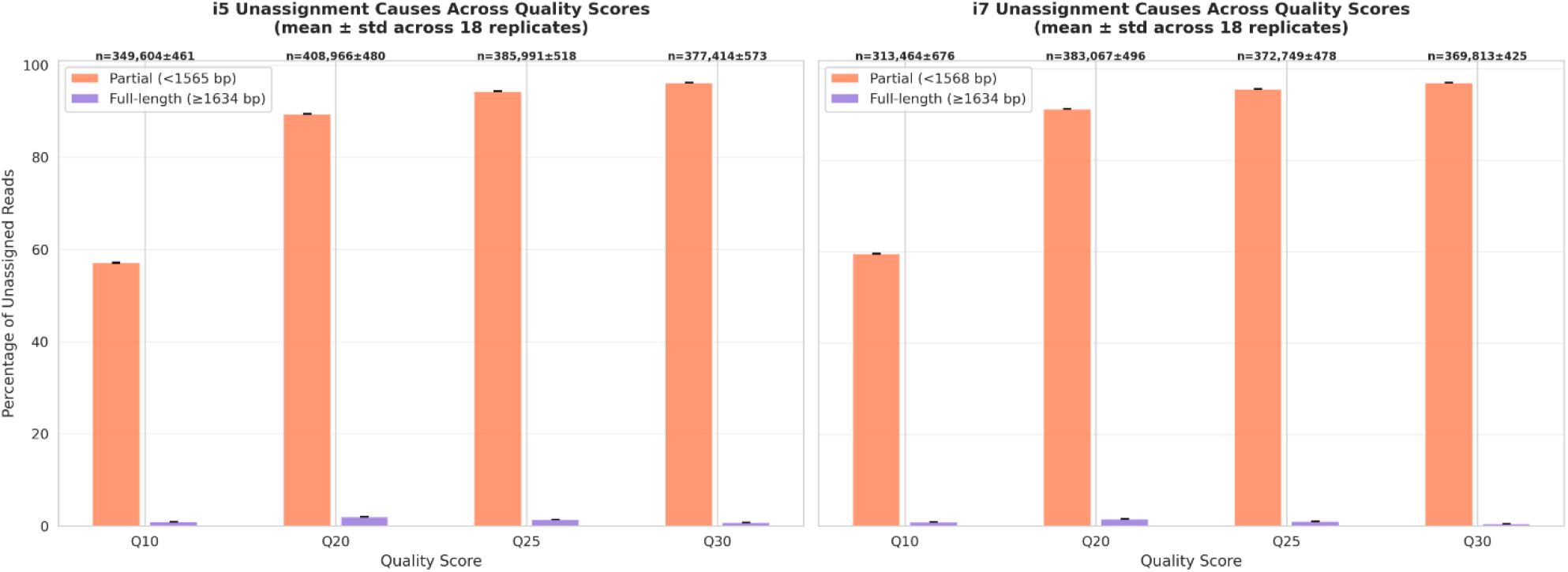
Causes of read unassignment across quality thresholds for the 96-sample design. Bars represent the percentage of unassigned reads categorized by sequence length: Partial (orange) and Full-length (purple). Results are shown for both i5 (left) and i7 (right) indexes. The total number of unassigned reads (n) is indicated above each bar. Data represents mean ± standard deviation across 18 replicates.

